# *Smg5* is required for multiple nonsense-mediated mRNA decay pathways in *Drosophila*

**DOI:** 10.1101/219725

**Authors:** Jonathan O. Nelson, Dominique Förster, Kimberly A. Frizzell, Stefan Luschnig, Mark M. Metzstein

## Abstract

The nonsense-mediated mRNA decay (NMD) pathway is a cellular quality control and post-transcriptional gene regulatory mechanism and is essential for viability in most multicellular organisms. A complex of proteins has been identified to be required for NMD function to occur, however the individual contribution of each of these factors to the NMD process is not well understood. Central to the NMD process are two proteins Upf1 (SMG-2) and Upf2 (SMG-3), which are found in all eukaryotes and are absolutely required for NMD in all organisms in which it has been examined. The other known NMD factors, Smg1, Smg5, Smg6, and Smg7 are more variable in their presence in different orders of organisms, and are thought to have a more regulatory role. Here we present the first genetic analysis of the NMD factor *Smg5* in *Drosophila*. Surprisingly, we find that unlike the other analyzed *Smg* genes in this organism, *Smg5* is essential for NMD activity. We found this is due at least in part to a role for Smg5 in the activity of two separable NMD-target decay mechanisms: endonucleolytic cleavage and 5′-to-3′ exonucleolytic decay. Redundancy between these degradation pathways explains why some *Drosophila* NMD genes are not required for all NMD-pathway activity. We also found that while the NMD component *Smg1* has only a minimal role in *Drosophila* NMD during normal conditions, it becomes essential when NMD activity is compromised by partial loss of *Smg5* function. Our findings suggest that not all NMD complex components are required for NMD function at all times, but instead are utilized in a context dependent manner *in vivo*.

## INTRODUCTION

Eukaryotic cells utilize a number of pathways to maintain error-free translation so as to preserve the fidelity of protein function (Adjibade and Mazroui 2014). Nonsense-mediated mRNA decay (NMD) is one such pathway, which prevents the translation of potentially harmful truncated proteins by recognizing and destroying mRNAs that contain erroneous premature-termination codons (PTCs) (Celik *et al*. 2015). In addition to this cellular quality control function, NMD degrades many endogenous wild-type mRNAs as a mechanism of post-transcriptional gene regulation (Peccarelli and Kebaara 2014).

While the phenomenon of NMD has been well characterized for several decades, the mechanisms initiating target recognition and degradation are still not well understood and it remains unclear if all the factors required for NMD activity have even been identified. Genes required for NMD were first found by genetic screens in yeast and *C. elegans*, which led to the identification of seven proteins required for NMD (Hodgkin *et al*. 1989; Leeds *et al*. 1991; 1992; Cali *et al*. 1999). Three of these genes, Upf1, Upf2, and Upf3, are present in every eukaryote examined, while the other four, Smg1, Smg5, Smg6, and Smg7, have variable presence across species (Siwaszek *et al*. 2014). In the absence of any one of these factors, PTC-containing mRNAs and endogenous targets are not efficiently degraded and instead accumulate in the cell (Gatfield *et al*. 2003; Rehwinkel *et al*. 2005). The molecular identities and biochemical characterization of the individual NMD genes have revealed clues about their roles in the NMD pathway. Upf1 is an ATP-dependent RNA helicase, and this activity is required for NMD (Czaplinski *et al*. 1995; Weng *et al*. 1996a; b). Upf3 binds mRNAs both directly and through an interaction with the exon-exon junction complex (EJC) (Gehring *et al*. 2003). Upf2 binds both Upf1 and Upf3, bridging an interaction between these two factors (He *et al*. 1997; Lykke-Andersen *et al*. 2000), helping stabilize Upf1-mRNA interactions. *Smg1* encodes a PIKK-like kinase that can phosphorylate Upf1. Loss of *Smg1* leads to reduced phospho-Upf1 in all organisms examined (Page *et al*. 1999; Yamashita *et al*. 2001; Grimson *et al*. 2004). In contrast, Upf1 is hyper-phosphorylated in *C. elegans smg-5, smg-6*, or *smg-7* mutants in a Smg1-dependent manner (Page *et al*. 1999), and RNAi inhibition of *Smg5, Smg6*, or *Smg7* in mammalian cells also results in Upf1 hyper-phosphorylation (Okada-Katsuhata *et al*. 2012). The finding that loss of any of the *Smg* genes reduce the efficiency of the NMD pathway even though they result in opposite effects on the Upf1 phosphorylation state has led to the concept that a cycle of Upf1 phosphorylation/dephosphorylation is a critical aspect of the NMD process (Ohnishi *et al*. 2003). The importance of Upf1 phosphorylation may be due to the 14-3-3-like domain found in Smg5, Smg6, and Smg7 proteins (Fukuhara *et al*. 2005). This domain binds phosphorylated residues, suggesting that Upf1 phosphorylation by Smg1 initiates binding of these factors to an NMD complex (Ohnishi *et al*. 2003). Smg6 is an endonuclease that cleaves targeted mRNAs near the PTC site (Gatfield and Izaurralde 2004; Huntzinger *et al*. 2008), suggesting that Smg6 binding to Upf1 is likely required for degradation of NMD targets. The functions of Smg5 and Smg7 are less clear, but a complex of Smg5 and Smg7 has been shown to bind a subunit of the PP2A phosphatase, suggesting Upf1 dephosphorylation may be mediated by these factors, likely after Smg6-mediated cleavage occurs (Anders *et al*. 2003; Ohnishi *et al*. 2003). Overall, these findings have led to a model in which Upf1 phosphorylation is critical for the NMD pathway, with Smg1 being required to phosphorylate Upf1 to recruit Smg6 and initiate NMD target cleavage, and Smg5 and Smg7 being required to dephosphorylate Upf1 to promote complex disassembly and recycling to new target mRNAs.

However, arguing against this model, recent studies dissecting the binding of Smg5, Smg6, and Smg7 to Upf1 suggest that Upf1 phosphorylation by Smg1 may not be key for normal NMD activity. It has been demonstrated that Smg5, Smg6, and Smg7 can bind Upf1 in the absence of Smg1, and indeed Smg6 binds Upf1 through a nonphosphorylated domain in the protein, indicating that Upf1 phosphorylation is not required for complex assembly (Nicholson *et al*. 2014; Chakrabarti *et al*. 2014). Additionally, Upf1 hyper-phosphorylation has been shown to mitigate the effects of reduced Smg5, Smg6, or Smg7 function (Durand *et al*. 2016). These findings suggest a revised model in which Smg5, Smg6, and Smg7 all contribute to the initiation of NMD target degradation independent of Upf1 phosphorylation, but when NMD activity is inefficient, Smg1 phosphorylates Upf1 to enhance the binding of Smg5 and Smg7, thus increasing NMD efficiency (Durand *et al*. 2016). Supporting this model, Smg5 and Smg7 have been shown in mammalian cell culture to interact indirectly with both decapping and deadenylation complexes (Cho *et al*. 2013; Loh *et al*. 2013), and thus may promote exonucleolytic degradation of NMD targets. Indeed, both endonucleolytic and exonucleolytic degradation products of endogenous NMD targets can be detected in mammalian cells (Lykke-Andersen *et al*. 2014; Schmidt *et al*. 2015; Colombo *et al*. 2017; Ottens *et al*. 2017). However, in part because experiments have been primarily performed in divergent cell lines, using different methods of gene manipulation and mostly studying transfected NMD target genes, it is unclear to what extent phosphorylation-independent Upf1-binding and the recruitment of decapping and deadenylation complexes occur during normal NMD activity *in vivo*.

NMD is required for viability in most complex organisms, including plants, *Drosophila*, zebrafish, and mice (Medghalchi *et al*. 2001; Yoine *et al*. 2006; Arciga-Reyes *et al*. 2006; Metzstein and Krasnow 2006; Weischenfeldt *et al*. 2008; Kerényi *et al*. 2008; Wittkopp *et al*. 2009; Li *et al*. 2015). *Drosophila* lacking *Upf1* or *Upf2* die during early larval stages, with no animals surviving to adulthood (Metzstein and Krasnow 2006; Chapin *et al*. 2014). However, *Drosophila* lacking *Upf3, Smg1*, or *Smg6* can survive to adulthood (Chen *et al*. 2005; Metzstein and Krasnow 2006; Avery *et al*. 2011; Frizzell *et al*. 2012). The viability of *Upf3, Smg1*, and *Smg6* mutants suggests that these animals have sufficient NMD activity to survive to adulthood, and indeed, these mutants display significant residual NMD activity. In particular, *Smg1* mutants show only a very small reduction in NMD activity (Chen *et al*. 2005; Metzstein and Krasnow 2006). *Smg5* is the only known *Drosophila* NMD gene for which loss-of-function mutations are yet to be described (the *Drosophila melanogaster* genome does not contain a *Smg7* orthologue (Chiu *et al*. 2003; Gatfield *et al*. 2003). Here we describe the first analysis of *Drosophila Smg5* mutants, and discover that *Smg5* is essential for NMD activity in this organism. By performing double-mutant analysis of NMD genes, we have found that Smg1 becomes essential for NMD when Smg5 function is compromised, and that Smg5 functions in a Smg6-independent degradation pathway *in vivo*. Our findings are consistent with the model that Smg1-mediated phosphorylation is only required under conditions of abnormal NMD progression, and that *Drosophila* utilize multiple independent mechanisms to initiate NMD target degradation.

## MATERIALS AND METHODS

**Fly genetics:** *Drosophila melanogaster* stocks were raised on standard cornmeal/dextrose food at 25°. The NMD mutant allele *Smg1^32AP^* (Metzstein and Krasnow 2006; Frizzell *et al*. 2012) is on a *y w FRT^19A^* chromosome and *Smg6^292^* (Frizzell *et al*. 2012) is on an *FRT^82B^* chromosome balanced over *TM6B, P{Dfd-EYFP} Sb^1^ ca^1^* (Le *et al*. 2006). All *Smg5* alleles are on *FRT^40A^* chromosomes and balanced over *CyO, P{Dfd-EYFP}* (Le *et al*. 2006). Other alleles used were *Gadd45^F17^* (Nelson *et al*. 2016), *pcm^14^* (Waldron *et al*. 2015), *Upf2^25G^* (Metzstein and Krasnow 2006), and *DHR78^3^* (Fisk and Thummel 1998). *y w FRT^19A^* was used as a control chromosome for all experiments for genes located on the X-chromosome (*Smg1* and *pcm* alleles). *FRT^40A^* or *FRT^82B^* were respectively used as the control chromosome for the experiments using *Smg5* or *Smg6* alleles. For viability tests, animals containing mutant alleles over a balancer were mated for three days, and offspring were collected for 10 days, beginning 10 days after mating was initiated. All offspring were scored, and percent expected viable was determined by the ratio of balancer negative animals to balancer positive animals.

**SV40 3′UTR constructs:** Deletion constructs of the SV40 3′UTR were made by one or two round PCR amplification with Phusion polymerase (NEB) using a pUASTeGFP::SV40 3′UTR plasmid as a template (Metzstein and Krasnow 2006). Amplicons were used to replace the full-length SV40 3′ UTR in a pUAST-eGFP plasmid modified to contain an *attB* site-specific recombination site (a gift from Jyoti Misra and Carl Thummel). All constructs were verified by sequencing and sent to Genetic Services, Inc (Cambridge, MA) for injection and site-specific integration into the *attP16* (2nd chromosome) strain (Venken *et al*. 2006). Expression levels of the modified SV40 3′UTR constructs were measured by mating transgenic males to *y w FRT^19A^; e22c-GAL4, UAS-nlsDsRed2::SV40 3′UTR/CyO* and *y w FRT^19A^ Upf2^25G^; e22c-GAL4, UAS-nlsDsRed2::SV40 3′UTR/CyO* females. Wandering third instar larvae expressing both DsRed and GFP were imaged using a Leica MZ16 fluorescence stereomicroscope.

**Screens for NMD-defective alleles:** The codon changes of all *Smg5* alleles can be found in **supplemental table 1**. For the mosaic genetic screen, males with an isogenized *FRT^40A^* 2^nd^ chromosome were starved for 8 hours, and then fed on sucrose containing 1% ethyl methanesulfonate overnight. Mutagenized males were then mated with *FRT^40A^*; *P{da-GAL4 w^+^} P{UAS-FLP} P{UAS-eGFP::SV40 3′UTR}* females (Wodarz *et al*. 1995), and F1 wandering L3 larvae were collected in glycerol and scored for mosaic enhanced GFP fluorescence using a Leica MZ 16F microscope equipped with epifluorescent illumination. Mutant mosaic animals were cleaned in PBS and placed in vials with food to continue development. After eclosion, candidate mutant lines were established and retested to confirm an NMD defect. Candidate alleles were tested for complementation with *Df(2L)BSC345* (Cook *et al*. 2012), which deletes the *Smg5* locus, and lines that failed to complement this deficiency for lethality or fluorescence enhancement were balanced over *CyO, P{Dfd:eYFP w^+^}* (Le *et al*. 2006).

The screen for embryos with enhanced reporter expression is described in Förster *et al*. (2010). Briefly, flies carrying NMD-sensitive UAS-GFP and UAS-Verm-mRFP reporters expressed in the tracheal system were mutagenized with EMS and F2 lines were established. F3s embryos were examined for reporter expression in tracheal cells.

**RNA isolation and quantification:** For qRT-PCR analyses, we collected five to ten larvae from 0-4 h after the L2-L3 molt and froze them in liquid nitrogen. We isolated total RNA using TRIzol reagent (Invitrogen) and Phase-Lock tubes (5-Prime), and the RNeasy mini kit (QIAGEN). We used on-column RNase-free DNase treatment (QIAGEN) to reduce genomic contamination. We determined RNA concentration by spectrophotometer and normalized concentration for reverse transcription. For reverse transcription, we used random decamers and MMLV8 reverse transcriptase (Retroscript Kit, Ambion). We performed qRT-PCR analysis using the SYBR Green qPCR Supermix (Bio-Rad) and the Bio-Rad iCycler thermocycler. All experimental reactions were performed using three technical replicates and a minimum of three biological replicates per condition, and the expression level of all experimental assays was normalized to *RpL32* mRNA expression. For all qRT-PCR analyses we also measured samples that had been made without reverse transcriptase to ensure that signal was not due to contamination with genomic DNA.

Primer sequences used were RpL32_1 (ATGCTAAGCTGTCGCACAAA), RpL32_2 (CGATGTTGGGGCATCAGATAC), Gadd45_5′_1 (CATCAACGTGCTCTCCAAGTC), Gadd45_5′_2 (CGTAGATGTCGTTCTCGTAGC), Gadd45_3′_1 (ACAGCCAGATGTCACAGAATT), and Gadd45_3′_2 (CCAGCAACTGGTTTCCATTAG). All *Gadd45* qPCR analysis was done using the Gadd45_5′ primer pair, unless otherwise noted.

**Analysis of *dHR78^3^* PTC allele stability:** We collected adult *Smg5^+/G115^* or *Smg5^C391/G115^* males that were also heterozygous for *dHR78^3^*, a PTC-containing allele that has lower expression than the wild-type allele and is stabilized in *Upf2^25G^* mutants, and thus is presumably degraded by NMD (Fisk and Thummel 1998; Nelson *et al*. 2016). At least three biological replicates were collected for each condition. We isolated RNA and generated cDNA, as described above, and used this cDNA as a template for PCR amplification of the *dHR78* transcript with the DRH78_F3 **/** DHR78_R3 primers (TGGGGCTTATTCAGAGTTCG / ATTAATGCTGGCCACACTCC), which flank the nonsense mutation. To compare the relative abundance of the *dHR78^3^* allele to the wild-type allele, PCR products were Sanger sequenced, and the relative peak intensity for a thymine (*dHR78^3^* allele) compared to a cytosine (wild-type allele) at nucleotide 1063.

**Lethal phase and larval development analysis:** For lethal phase and larval development analysis, first-instar larvae were collected 20-24 hours after egg lay. Every 24 hours, animals were examined to record their developmental stage and transferred to fresh food. Larval stage was determined based on physical characteristics of the mouth hooks. Once animals entered pupariation, pupae were transferred to vials and scored for eclosion five days later.

**Statistical Analysis:** All viability assay figures represent the proportion of animals of the indicated genotypes that survive to adulthood; error bars for these figures represent the 95% confidence interval of the binomial distribution, and the Test of Equal or Given Proportions was used to determine significance difference in these proportions between genotypes. For each individual experiment, conditions were compared directly to the control, so no p-value correction was applied. All other figures represent the mean value of multiple replicates and display error bars representing ± 2 SEM. For tests between two variable measures, a two-sided paired Student’s t-test was used to determine significance difference between mean value data. Those qPCR experiments that compared a condition to the control, which was set to a constant of 1, were performed with a one-sided Student’s t-test.

**Data Availability:** *Drosophila* strains are available upon request. All data is presented within the figures.

## RESULTS

**Isolation of *Smg5* mutant alleles:** To identify *Drosophila Smg5* mutant alleles, we used two different genetic screens. First, we performed an EMS-mutagenesis screen in mosaic animals expressing an NMD-sensitive GFP reporter, a method similar to one we previously used to recover *Smg6* alleles (Frizzell *et al*. 2012). This reporter expresses GFP from a pUAST construct (Brand and Perrimon 1993) bearing a UAS promoter and an NMD-sensitive *SV40* 3′UTR (Metzstein and Krasnow 2006). We generated mosaics using the *da-GAL4* driver to ubiquitously express FLP-recombinase **(Figure 1A)**. Individual homozygous mutant cells with defective NMD activity show increased reporter expression and GFP fluorescence **(Figure 1A)**. The mosaic enhanced fluorescence phenotype was easy to distinguish in late L3 larvae **(Figure 1B)**, and mosaic animals remain viable and fertile, so even lethal alleles can be recovered from individual mutants. An added benefit of this approach is that by mutagenizing animals that have an *FRT* site located near the centromere on the left arm of the second chromosome (*FRT^40A^*), we could specifically isolate mutations only on this chromosome arm. Since *Smg5* is located on the left arm of the second chromosome, mutations identified from the screen would likely include *Smg5* alleles. Using this approach, we screened 12,554 larvae and identified three mutants with mosaic enhancement of GFP fluorescence **(Figure 1C)**. We found each of these three mutants were homozygous lethal. We crossed each allele to a deficiency that deletes *Smg5* and found that all three failed to complement for lethality, suggesting that they had mutations in *Smg5*.

**Figure 1.**
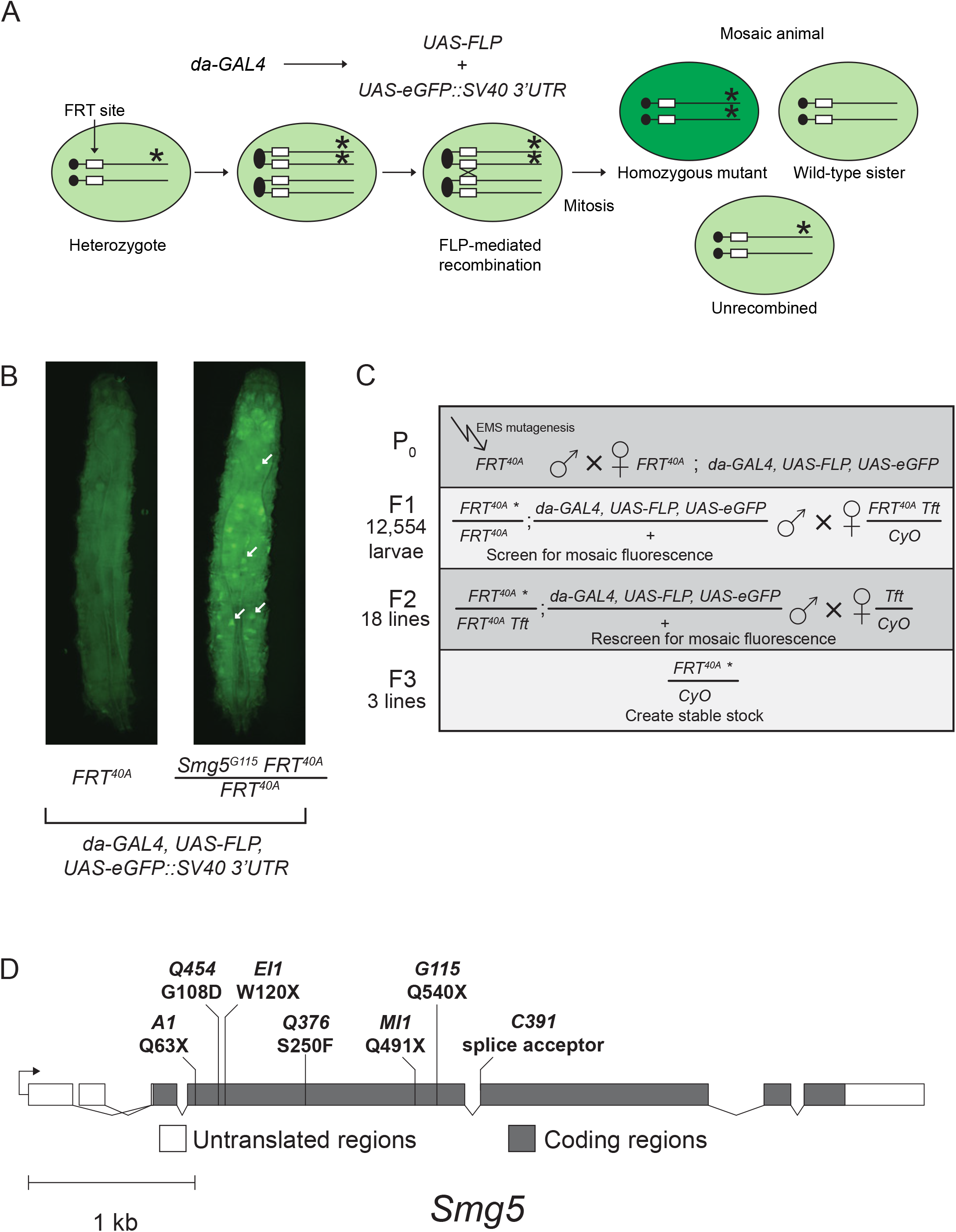
Mosaic screen for novel NMD-defective mutations on the 2^nd^ chromosome identifies *Smg5* alleles. (**A**) Scheme to generate mosaics and detect mutants with defective NMD. The GAL4 transcription factor is ubiquitously expressed under a *daughterless* (*da*) promoter and activates transcription of *FLP* recombinase and the NMD-sensitive *eGFP::SV40 3′UTR* fluorescent reporter, both under UAS control. The reporter mRNA is usually degraded by NMD, and thus cells lacking NMD activity due to a homozygous mutation in a gene required for NMD activity show increased green fluorescence. (**B**) Example of mosaic GFP-reporter fluorescence phenotype detected in our screen. Late L3 larvae expressing the NMD sensitive *eGFP::SV40 3′UTR* fluorescent reporter in animals with a wild-type *FRT^40A^* chromosome (left) or an *FRT^40A^ Smg5^G115^* chromosome (right). Individual homozygous mutant cells with increased GFP fluorescence in the *Smg5* mutant animal are indicated by white arrows. Overall increased fluorescence is due to other out of focus mutant cells. Dorsal view; anterior at top. **(C)** Scheme for recovering mutations identified in the screen. The total number of candidate mutants scored in each generation is shown on the left side of each row. Genotypes on the left in the F1 and F2 generations are the offspring from the previous mating. (**D**) Molecular identity of isolated *Smg5* mutations. Four alleles (*A1, EI1, MI1*, and *G115*) are nonsense mutations. *C391* is a mutation in a splice acceptor site. *Q454* and *Q376* are missense mutations. The codon changes of all *Smg5* alleles are listed in **Supplemental Table 1**.

Our second screen was of animals expressing a *GFP::SV40 3′UTR* reporter in the embryonic tracheal system (Förster *et al*. 2010). This screen identified four mutants that showed increased fluorescence **(Figure 1D, Supplemental Figure 1)**. All four of these alleles failed to complement a *Smg5* deficiency using increased fluorescence signal as an assay (data not shown), indicating they contained mutations in *Smg5*. As expected for mutations disrupting NMD-pathway function (Metzstein and Krasnow 2006), the increase in fluorescence was independent of the fluorescent reporter examined, with both GFP and mRFP showing similar increases in expression in a homozygous mutant background **(Supplemental Figure 1F)**.

Finally, sequencing of the *Smg5* locus in the seven candidate lines revealed they all contained mutations in *Smg5*, including nonsense mutations (*G115, A1, EI1*, and *MI1*), an altered splice acceptor site (*C391*), and missense mutations in highly conserved alpha-helices of the *Smg5* 14-3-3-like domain (*Q454* and *Q376*) (Fukuhara *et al*. 2005) **(Figure 1D; Supplemental Table 1)**.

***Smg5* is an essential NMD factor in *Drosophila:*** *Drosophila* lacking any functional NMD activity, such as *Upf1* and *Upf2* null mutants, fail to develop to adulthood, dying primarily during early larval stages (Chapin *et al*. 2014). We found that the *Smg5* nonsense alleles *A1, EI1, MI1*, and *G115*, and splice acceptor site allele *C391* all failed to survive to adulthood when over a deficiency that deletes *Smg5*, or as *Smg5^C391/G115^* transheterozygotes **(Figure 2A)**. The lethality of these alleles combined with their molecular aberrations suggested that they are complete loss-of-function mutations. We found that *Smg5^C391/G115^* mutants have developmental delays, with *Smg5* mutants spending almost twice as long in larval stages as control animals **(Supplemental Figure 2A)**, and most *Smg5^C391/G115^* mutants die during pupariation **(Supplemental Figure 2B)**. This developmental delay and lethal phase is similar to, but somewhat weaker than, the developmental defects of null *Upf1* and *Upf2* mutants (Chapin *et al*. 2014). Conversely, the missense alleles *Q454* and *Q376* were viable over the deficiency and each other **(Figure 2A)**, suggesting that these are likely hypomorphic alleles.

**Figure 2.**
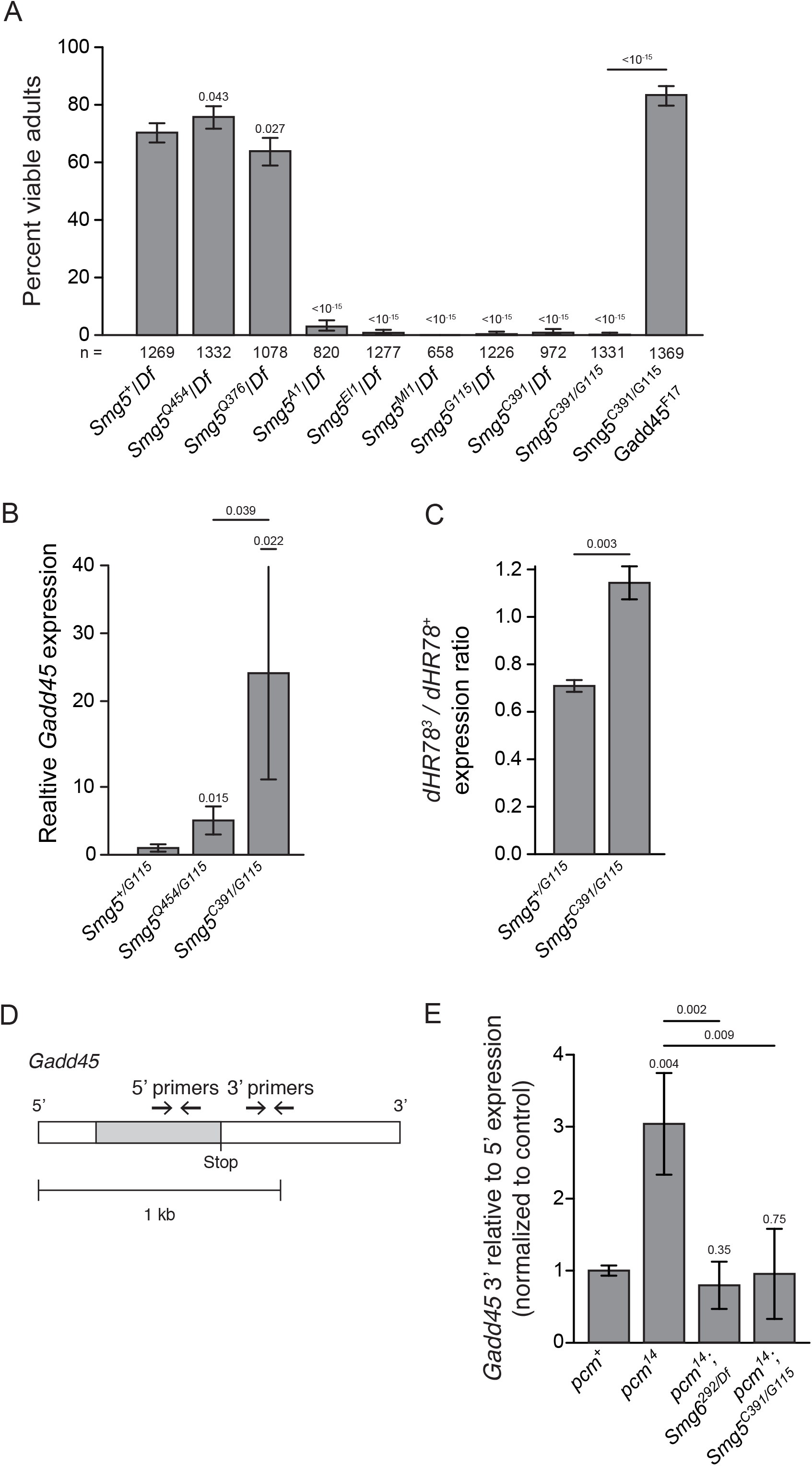
*Smg5* is required for viability and NMD activity in *Drosophila*. (**A**) Adult viability of *Smg5* mutant alleles *trans*-heterozygous to either a deficiency removing the *Smg5* locus (*Df*) or other *Smg5* mutant alleles. Error bars represent 95% confidence interval of the binomial distribution. p-value listed compared to *Smg5^+^* / *Df* condition or between indicated conditions determined by the test of equal or given proportions. n = total number of animals scored. (**B**) Expression of the endogenous NMD target *Gadd45*, as measured by qRT-PCR. Error bars represent 2 SEM. p-value listed for each condition compared to controls or between indicated conditions, determined by two-sided Student’s t-test. n ≥ 3 for all conditions. (**C**) Relative abundance of PTC-containing *dHR78^3^* allele (Fisk and Thummel 1998) mRNA compared to wild-type *dHR78* allele mRNA in animals heterozygous for *dHR78^3^* in each indicated genotype. Error bars represent 2 SEM. p-value listed for each condition compared to + / *Smg5^G115^*, determined by two-sided Student’s t-test. n = 3 for all conditions. **(D)** Diagram of the endogenous NMD target *Gadd45* transcript and 5′ and 3′ qRT-PCR primer pairs. Open boxes indicate UTRs; grey boxes indicate coding regions. 5′ primer pair is located 5′ to the stop codon and the 3′ primer pair is 3′ to the stop codon. **(E)** *Gadd45* expression measured with the 3′ primer pair relative to the 5′ primer pair. The 3′ region is preferentially stabilized in *pcm^14^* mutants. This preferential stabilization is lost when either *Smg6* or *Smg5* are lost. Error bars represent 2 SEM. p-value listed for each condition compared to the *pcm^+^* condition or between indicated conditions, determined by two-sided Student’s t-test. n ≥ 3 for all conditions.

Lethal mutations in *Drosophila* NMD genes generally have severe defects in NMD function, as measured by increased expression of endogenous NMD targets (Metzstein and Krasnow 2006; Avery *et al*. 2011; Frizzell *et al*. 2012). To test if lethal *Smg5* mutant alleles also have strong defects in NMD activity, we used qRT-PCR to measure the expression of the endogenous NMD target *Gadd45* (Chapin *et al*. 2014; Nelson *et al*. 2016). Since *Gadd45* is directly targeted by NMD, the amount of *Gadd45* mRNA in mutants serves as a measure of the decrease in NMD activity. We measured *Gadd45* mRNA levels in early third instar larvae and found that *Smg5^C391^/G115* mutants had a large increase in *Gadd45* mRNA expression. In contrast, viable *Smg5^Q454/G115^* mutants showed a much smaller increase in *Gadd45* levels **(Figure 2B)**. Increased *Gadd45* expression is a major factor contributing to the death of *Upf1* and *Upf2* mutants, and loss of *Gadd45* can suppress *Upf1* and *Upf2* mutant lethality (Nelson *et al*. 2016). We found that loss of *Gadd45* also suppresses the lethality of *Smg5^C^391/G115* mutants **(Figure 2A)**, indicating that these animals are dying due to a similar loss of NMD function as *Upf1* or *Upf2* mutants. These results strongly suggest that *Smg5* mutant lethality is specifically due to a loss of NMD activity, and not due to loss of any NMD-independent Smg5 function.

***Smg5* null mutants lack most, if not all, detectable NMD activity:** To directly test if *Drosophila Smg5* mutants have any residual NMD activity, we measured the relative stability of PTC-containing mRNAs in *Smg5* mutants. We found that *Smg5^C391/G115^* mutants fully stabilized the expression of the PTC-containing *dHR78^3^* mRNA relative to the expression of wild-type *dHR78* mRNA **(Figure 2C)**, indicating NMD-mediated degradation of PTC-containing mRNA is absent. Since Smg6-mediated cleavage is a known mechanism for degradation of NMD targets in *Drosophila*, we tested if *Smg5* mutants still retain this endonuclease activity. NMD-target cleavage can be observed through measuring the relative abundance of NMD-target mRNA fragments 5′ to the stop codon in relation to fragments 3′ to the stop codon **(Figure 2D)** in animals lacking the only cytoplasmic 5′-to-3′ exonuclease Xrn1, which is encoded by the gene *pacman* (*pcm*) (Till *et al*. 1998). Null *pcm* mutants have no 5′-to-3′ exonuclease activity (Waldron *et al*. 2015), and thus mRNAs cleaved by Smg6 near the stop codon show increased abundance of the 3′ cleavage fragment compared to the 5′ fragment (Nelson *et al*. 2016). We found this bias to be lost in the absence of *Smg6* **(Figure 2E)**, confirming that it is caused by Smg6 endonuclease activity. Interestingly, we found that the preferential stabilization of the 3′ fragment is also lost in double mutants of the null alleles of *pcm* and *Smg5* **(Figure 2E)**, revealing that Smg5 is required for Smg6 endonuclease activity. These combined results indicate that *Drosophila Smg5* mutants lack any NMD activity.

As an additional gauge of NMD activity in *Smg5* mutants, we directly measured fluorescence levels of NMD-sensitive reporters in homozygous mutant embryos **(Figure 3)**. We found that homozygous *Smg5^C391^* embryos exhibited ∼5-fold increase in fluorescent signal compared to *Smg5^+^* embryos, comparable to the increase in GFP mRNA levels observed in the strongest previously measured NMD mutant, *Upf2^25G^* (Metzstein and Krasnow 2006). As expected, embryos homozygous for the hypomorphic allele *Smg5^Q454^* showed a smaller increase in fluorescent signal, while the transheterozygous combination *Smg5^C391^/Q454* showed an intermediate signal, close to the *Smg5^Q454^* signal. To provide a direct comparison of the *Smg5* mutant fluorescence to other reporters, we generated a series of deletion constructs of the NMD-sensitive SV40 3′UTR and tested each construct for expression level and ability to be enhanced by loss of NMD activity **(Supplemental Figure 4)**. We found that, in general, the shorter the deletion construct the higher the absolute expression level and the less this expression was enhanced by loss of NMD, though for a given length of construct, the exact location of the deletion determined the degree of enhancement. These results are consistent with a model in which 3′UTR length is a major determinant of NMD sensitivity (Boehm *et al*. 2014), though there is likely a contribution of specific sequence elements in modulating sensitivity. Most importantly, we found the construct with greatest enhancement in expression increased fluorescence levels 5-fold compared to the full-length SV40 3′UTR, almost exactly the same as the increase observed in the *Smg5* null mutant background **(Figure 3)**. Hence, we conclude that loss of *Smg5* phenocopies loss of NMD sensitivity, suggesting *Smg5* is required for all NMD activity in embryos.

**Figure 3.**
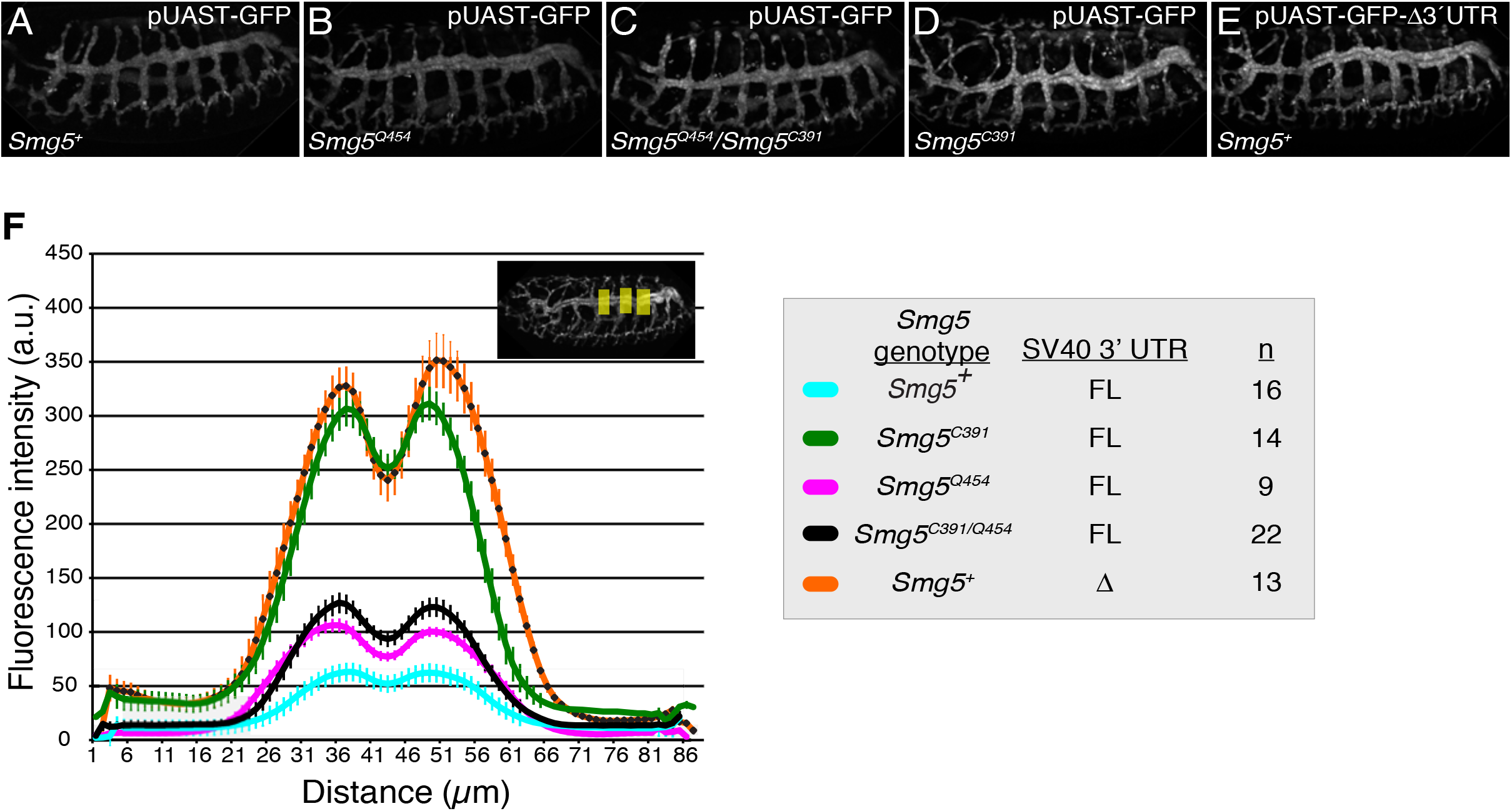
Loss of *Smg5* enhances NMD-sensitive reporter expression in embryos. **(AE)** embryos of indicated genotypes carrying GFP reporter transgene with full-length NMD-sensitive SV40 3′UTR (pUAST-GFP) or NMD-insensitive SV40 3′UTR deletion construct (pUAST-GFP-∆3´UTR). **(F)** Scan of GFP intensity averaged across three areas of embryo (shown in inset) in animals of the indicated genotype. FL, full-length SV40 3′UTR; ∆, SV40 3′UTR deletion construct; n, number of embryos scored.

**Figure 4.**
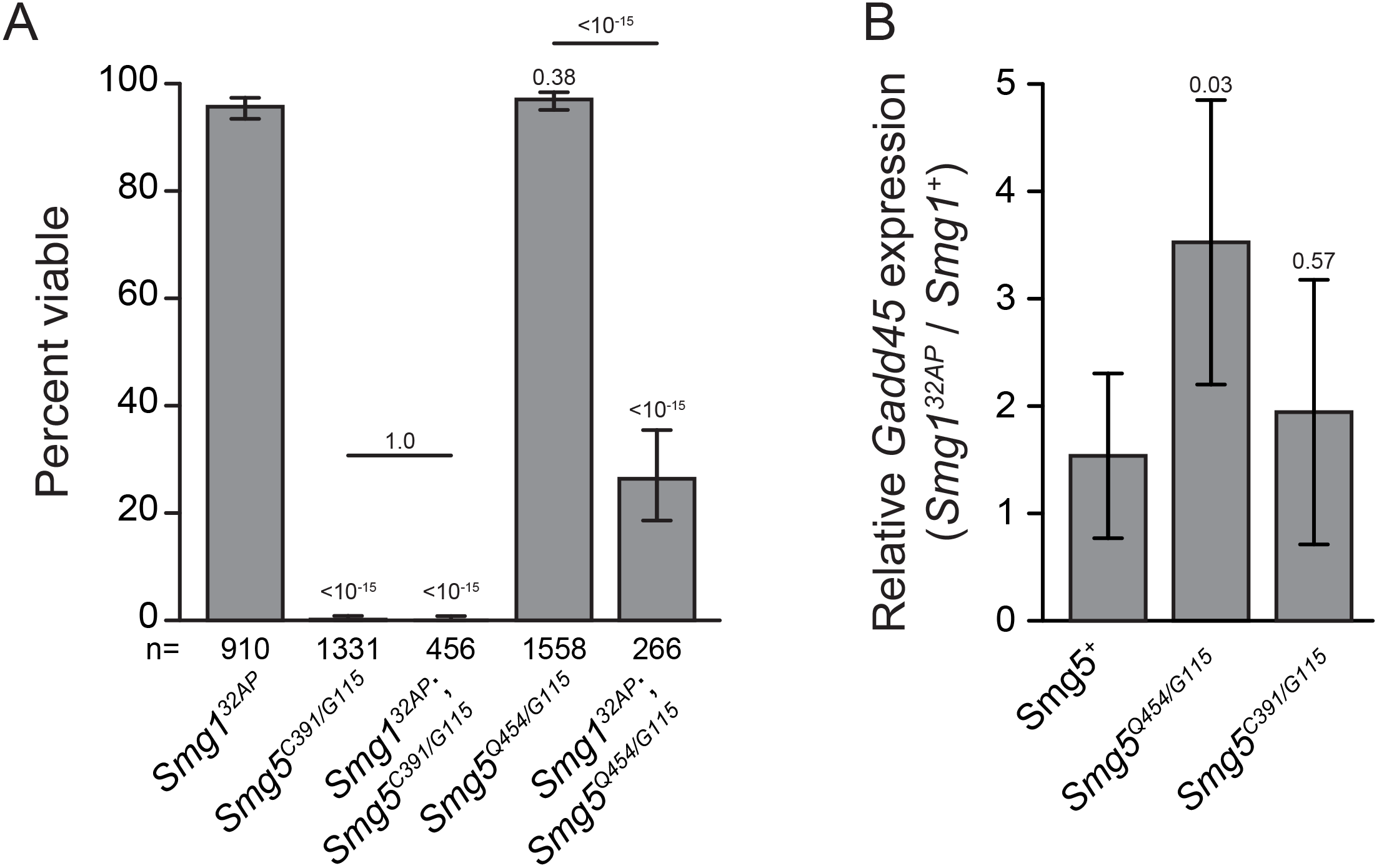
*Smg5* mutant lethality is not dependent on *Smg1*. (**A**) Adult viability of *Smg1* and *Smg5* hypomorph single and double mutants. Error bars represent 95% confidence interval of the binomial distribution. p-value listed for each condition compared to *Smg1^32AP^* or between indicated conditions determined by the test of equal or given proportions. n = total number of animals scored. **(B)** Relative expression of the endogenous NMD target *Gadd45* in *Smg1^32AP^* mutants compared to *Smg1^+^* controls in *Smg5^+^* and mutant backgrounds as measured by qRT-PCR. Error bars represent 2 SEM. p-value listed for each condition compared to *Smg5^+^* determined by two-sided Student’s t-test. n ≥ 3 for all conditions.

***Smg5* mutant lethality is not *Smg1*-dependent:** The phosphorylation of Upf1 by Smg1 has been proposed to be a critical step in the NMD process, at least in part by recruiting Smg6 to the NMD complex to initiate target degradation (Hug *et al*. 2016).

Dephosphorylation of Upf1 is thought to be mediated by Smg5, which interacts with the PP2A phosphatase; this activity may be required for complex disassembly after target degradation has been initiated (Ohnishi *et al*. 2003). This model thus proposes that the necessity of Smg5 for NMD activity requires Upf1 phosphorylation by Smg1, and thus it would be expected that *Smg1* mutants, which are fully viable and have robust NMD activity (Chen *et al*. 2005; Metzstein and Krasnow 2006; Frizzell *et al*. 2012), should suppress *Smg5* mutant lethality. In contrast to this prediction, we found that *Smg1*; *Smg5* double mutants were in fact no more viable than *Smg5* mutants **(Figure 4A)**, and *Smg1* mutants had no effect on the developmental delay or lethal stage of *Smg5* mutants **(Supplemental Figure 2A, B)**. These findings suggest that a failure to dephosphorylate Upf1 is not responsible for *Smg5* mutant lethality; however, the lethality of *Smg1*; *Smg5* double mutants may also be explained by unknown factors that phosphorylate Upf1 in the absence of Smg1.

If failure to dephosphorylate Upf1 causes lethality in both *Smg5* mutants and *Smg1; Smg5* double mutants, we would expect loss of *Smg1* to have no effect on the viability of hypomorphic *Smg5* mutants, since these alleles are viable **(Figure 2A)**, and so should have sufficient Upf1-dephosphorylation. Surprisingly, we found that double mutants for a *Smg1* null allele and a hypomorphic *Smg5* allele show significant lethality, even though each mutation on its own is viable **(Figure 4A)**. This result reveals that *Smg5* mutant lethality is not due to failure to dephosphorylate Upf1, but instead is consistent with an alternative proposed model that Upf1 phosphorylation by Smg1 is not required for NMD under normal conditions, but serves to enhance Smg6 and Smg5 efficiency upon stress conditions to reinforce NMD activity (Durand *et al*. 2016). In agreement with this model, we found that the relative increase in *Gadd45* expression upon loss of *Smg1* is greater in this *Smg5* hypomorphic background than in animals with functioning *Smg5* **(Figure 4B)**, indicating that Smg1 has a greater contribution to NMD activity when Smg5 functions inefficiently. Importantly, loss of *Smg1* has no greater impact on *Gadd45* expression in *Smg5* null mutants than in animals with functional *Smg5* **(Figure 4B)**, indicating that the compensatory Smg1 activity in *Smg5* hypomorphs is the enhancement of Smg5 function, rather than an increase in Smg5-independent decay activity. Together, these findings indicate that the requirement of Smg5 for NMD activity is independent of Smg1, but that Smg1 can enhance Smg5 activity when NMD function is compromised.

**Smg1 is not required for Smg6 activity:** Smg6 has been shown to bind Upf1 at residues phosphorylated by Smg1 (Fukuhara *et al*. 2005; Okada-Katsuhata *et al*. 2012), leading to the model that Smg1 is required for Smg6 complex entry and cleavage of NMD targets. However, *Drosophila Smg6* mutants have much stronger NMD defects than *Smg1* mutants (Frizzell *et al*. 2012), and Smg6 has recently been shown to be capable of binding non-phosphorylated Upf1 (Nicholson *et al*. 2014; Chakrabarti *et al*. 2014). These data suggest an alternative model in which Smg6 can cleave NMD targets even in the absence of Smg1 kinase activity. In support of this latter model, we found using our 3′ fragment stabilization assay that *Gadd45* mRNA is cleaved in the absence of *Smg1* just as efficiently as in animals with wild-type Smg1 **(Figure 5A)**. We also found the same relative increase in *Gadd45* mRNA levels upon loss of *Smg6* in animals with or without functional Smg1 **(Figure 5B)**. Consistent with this finding, double mutants between null *Smg1* and *Smg6* alleles do not have reduced viability or enhanced developmental delay compared to *Smg6* single mutants **(Figure 5C; Supplemental Figure 2C)**. Together, these data suggest that Smg1 does not contribute to NMD-independent Smg6 function, and that Smg1is not required for normal Smg6 activity *in vivo*.

**Figure 5.**
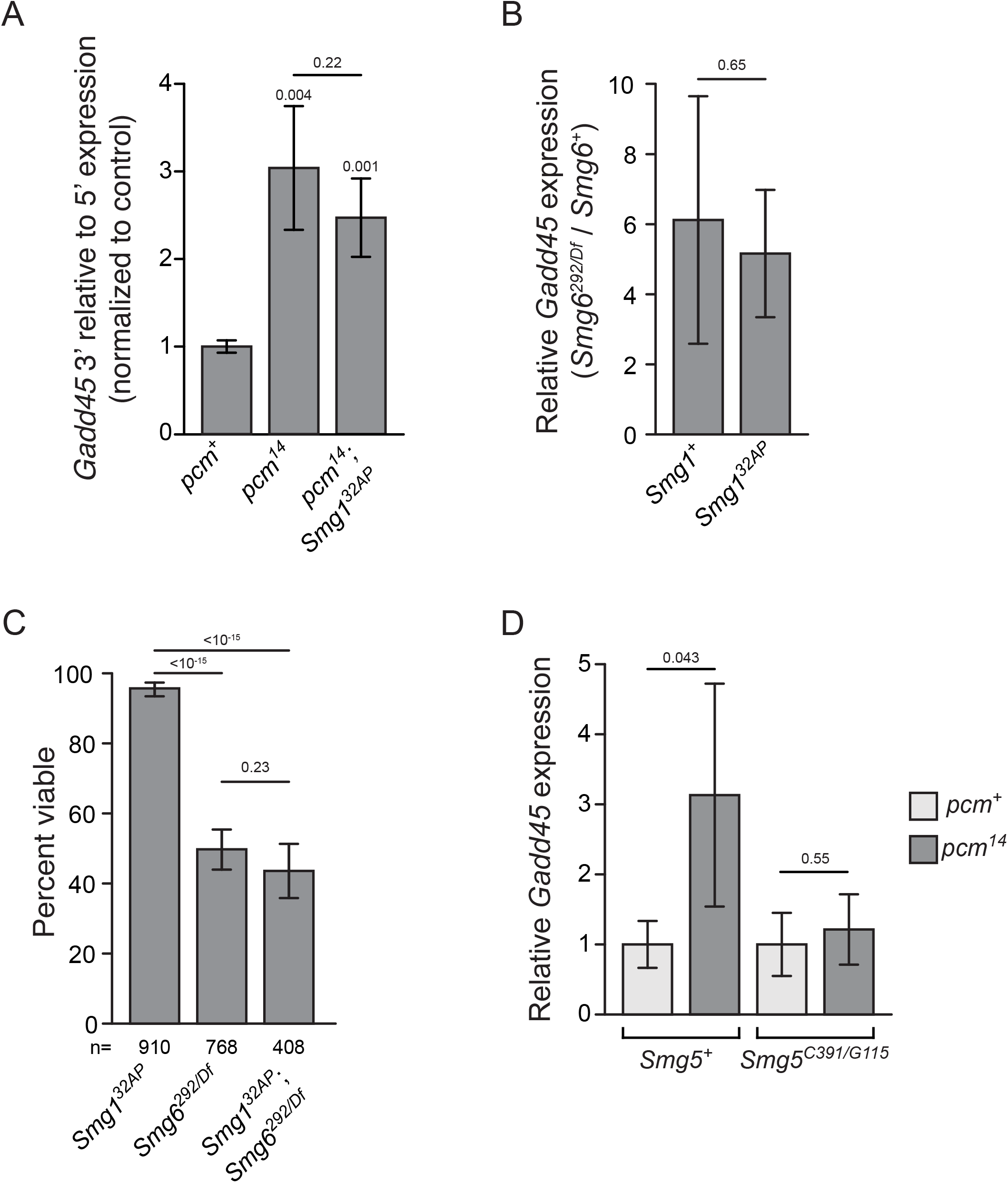
*Smg1* is not required for Smg6 activity and *Smg5* is required for exonucleolytic NMD activity. (**A**) *Gadd45* 3′ expression relative to 5′ expression in indicated genotypes, measured by qRT-PCR. The 3′ region is preferentially stabilized in *pcm^14^* mutants with or without functional *Smg1*. Error bars represent 2 SEM. p-value listed for each condition compared to *pcm^+^* or between indicated conditions determined by two-sided Student’s t-test. n ≥ 3 for all conditions. (**B**) Relative expression of the endogenous NMD target *Gadd45* measured by qRT-PCR using the 5′ primer pair in *Smg6^292/Df^* mutants compared to *Smg6^+^* controls in either *Smg1^+^* or *Smg1^32AP^* mutant backgrounds. Error bars represent 2 SEM. p-value between conditions determined by two-sided Student’s t-test. n ≥ 3 for all conditions. **(C)** Adult viability of *Smg1^32AP^* and *Smg6^292^/Df* null mutants, and *Smg1^32AP^; Smg6^292/Df^* double mutants. Error bars represent 95% confidence interval of the binomial distribution. p-value listed between indicated conditions determined by the test of equal or given proportions. n displays total number of animals scored. (**D**) Relative expression of the endogenous NMD target *Gadd45* in *pcm* mutants as measured by qRT-PCR. Expression is normalized to the *pcm^+^* condition in either a *Smg5^+^* or *Smg5^C391/G115^* condition. Error bars represent 2 SEM. p-value listed between indicated conditions determined by two-sided Student’s t-test. n ≥ 3 for all conditions.

***Smg5* is required for 5′-to-3′ Xrn1-mediated exonucleolytic degradation of NMD targets:** The stronger defects observed in *Smg5* mutants than *Smg6* mutants indicate that more than just Smg6 activity is disrupted in *Smg5* mutants. Smg5 has been shown in mammalian cell culture to interact indirectly with decapping and deadenylation complexes (Cho *et al*. 2013; Loh *et al*. 2013), however if either of these interactions occur during *in vivo* NMD targeting is unclear. Decapping of NMD targets is expected to lead to Xrn1-mediated 5′-to-3′ exonucleolytic degradation. While inhibition of decapping factors or Xrn1 in *Drosophila* cells was found to have no effect on the levels of a PTC-containing reporter detected by northern blot (Gatfield and Izaurralde 2004), we have identified an increase in *Gadd45* expression in null *Xrn1* mutants by qRT-PCR *in vivo* **(Figure 5D)** (Nelson *et al*. 2016). Although this increase could be due to NMD-independent Xrn1 activity, we found that there is no difference in *Gadd45* expression upon loss of *Xrn1* activity in a *Smg5* mutant background **(Figure 5D)**, indicating that Xrn1-mediated degradation of *Gadd45* mRNA requires Smg5 activity. Importantly, *Gadd45* expression was measured using the qRT-PCR primer pair 5′ to the *Gadd45* stop codon **(Figure 2D)**, implying that Xrn1 can only degrade these mRNAs after decapping. These findings suggest that Smg5 may be involved in promoting the decapping of NMD targets.

## DISCUSSION

The degradation of both specific normal and many kinds of erroneous mRNAs by the NMD pathway is a crucial gene regulatory mechanism and arose in the ancestors to all eukaryotes. While many factors required for NMD have been biochemically characterized, the individual contribution of each factor to the recognition and degradation of NMD targets is not fully understood, especially *in vivo* in complex organisms. Through our genetic analysis of *Smg5* in *Drosophila* and by examination of NMD-gene double mutants, we have found that NMD utilizes multiple mechanisms to promote target degradation *in vivo*. One of our main findings is that *Smg5* null mutants have as severe defects as either *Upf1* and *Upf2* null mutants, indicating that *Smg5* is a critical factor for promoting NMD-target recognition and/or decay. In support of this interpretation, we found that Smg5 is required for both Smg6-mediated endonucleolytic cleavage of NMD targets and a separate, Smg6-independent, decay process that at least partially requires Xrn1 5′-to-3′ exonuclease activity. Our findings are surprising, given that *Smg5* has primarily been thought to promote NMD complex recycling, but with only a secondary requirement to stimulate decay activity (Ohnishi *et al*. 2003). Instead, we propose that Smg5 is a critical NMD factor necessary for at least two, independent NMD degradation mechanisms.

In contrast to Smg5 having a critical role in target degradation, our data is less supportive for a Smg5 function in NMD complex recycling. The phosphorylation of Upf1 at multiple residues by Smg1 (Yamashita *et al*. 2001; Grimson *et al*. 2004) is thought to be required to initiate, or at least stimulate, NMD-mediated degradation (Anders *et al*. 2003; Ohnishi *et al*. 2003). Subsequent Smg5-mediated recruitment of the PP2A phosphatase to the NMD complex is thought to lead to Upf1 dephosphorylation to promote complex disassembly and recycling (Anders *et al*. 2003; Ohnishi *et al*. 2003). While Smg1, and thus Upf1 phosphorylation, does not seem to play a major role in NMD in *Drosophila* (Chen *et al*. 2005; Metzstein and Krasnow 2006), failure to dephosphorylate Upf1 could still be the cause of the strong NMD defect in *Drosophila Smg5* mutants. This model predicts that since *Smg5* mutant lethality is due to lack of Upf1 dephosphorylation, the loss of *Smg1* should be epistatic to the loss of *Smg5*, since Upf1 dephosphorylation would no longer be required in *Smg1* mutant animals. However, we found just the opposite: *Smg1* mutations do not suppress *Smg5* mutations at all, and *Smg1* mutants actually enhance the defect of *Smg5* hypomorphic alleles. These data are instead consistent with a more recently proposed model in which Upf1 phosphorylation is only required when the NMD process is “stalled” or otherwise becomes impaired

(Durand *et al*. 2016), such as occurs in hypomorphic *Smg5* mutants. Based on our findings, we propose that NMD functions under two states: non-phosphorylated Upf1, which requires Smg5 to stimulate both Smg6-mediated and Smg6-independent decay, and phosphorylated Upf1, which enhances an interaction of Smg6 with Upf1 and stimulates Smg6-mediated decay independently of Smg5. For most substrates under normal conditions the former mechanism predominates, with the later only occurring when the first process does not efficiently occur. Additionally, some specific substrates may require Smg1-stimulated NMD by default. For instance, while loss of Smg1 leads to barely detectable stabilization of a PTC-containing substrate (Chen *et al*. 2005; Metzstein and Krasnow 2006), there is a significant (>2-fold) increase in endogenous substrates, such as *Gadd45*, and reporters using the NMD-sensitive SV40 3′UTR (Metzstein and Krasnow 2006; Frizzell *et al*. 2012). The difference in targeting between these alternative substrates is not yet known, but there is indication of differential pathway usage also in mammalian cells (Chan *et al*. 2007; Ottens *et al*. 2017).

The stronger NMD defects observed in *Smg5* null mutants compared to *Smg6* null mutants suggest that Smg5 is required for a Smg6-independent decay activity. While the mechanism of this decay remains unclear, there is *in vitro* evidence that Smg5 can interact with the Dcp decapping complex (Cho *et al*. 2009; 2013; Loh *et al*. 2013), recruiting the complex to the NMD target and eventually leading to exonucleolytic decay initiated at the 5′ end of the message. Additionally, global analysis of NMD-decay intermediates suggests that the degradation of NMD substrates mainly occurs through Smg6-mediated endonucleolytic cleavage, but that substrates bypassing this decay process may go through an alternative decapping-mediated process (Lykke-Andersen *et al*. 2014). However, whether this alternative process is actually used *in vivo*, and whether it depends on Smg5 has not been genetically examined. By examining decay in *Xrn1 Smg5* double mutants we have not obtained such evidence. Xrn1 is expected to have a role in both Smg6-mediated endonucleolytic decay and 5′ decapping decay, but with measurably different effects. In Smg6-mediated decay, Xrn1 is expected to be required for the degradation of the RNA fragment 3′ to the cleavage site, and we and others have demonstrated such a requirement *in vivo* and in cell culture. However, in decapping-dependent decay, Xrn1 is expected to be required for the degradation of the entire mRNA, including sequence 5′ to the cleavage site. Here, we have shown that degradation of an NMD substrate 5′ to the predicted Smg6-cleavage site is in at least in part dependent on both Smg5 and Xrn1, indicating an *in vivo* role for Smg5-mediated decapping in NMD substrate degradation. The incomplete loss of NMD activity in *Smg6* mutants suggests that Smg6-indepdendent decay is sufficient to maintain most NMD activity. It is also possible that the preference for which decay mechanism degrades NMD targets may be different between individual NMD targets. Furthermore, the choice between decay mechanisms may differ in tissue-specific or developmental contexts. It will be important to parse the relative contribution of each decay pathway to the degradation of NMD targets to understand the mechanism underlying the bias in decay.

Here we performed the first double mutant analysis of multiple NMD factors, providing genetic evidence of the relative contribution of individual NMD genes. We also characterized the first *Drosophila Smg5* mutants, identifying that *Smg5* is critical for NMD function and viability, similar to *Upf1* and *Upf2*, and providing the first genetic evidence for an essential role of *Smg5* function in a model system. Our findings suggest that NMD utilizes multiple branched decay mechanisms to destroy its targets. All of these pathways depend on *Smg5*, indicating that *Smg5* plays more fundamental roles in NMD than has previously been appreciated. More closely characterizing the molecular mechanisms of Smg5 function in NMD may reveal novel key features of NMD activity that have thus far escaped detection.

## Acknowledgements

We thank x for critical comments on the manuscript. We are very grateful to Sarah Newbury for providing fly stocks prior to publication. Fly stocks were obtained from the Bloomington *Drosophila* Stock Center. KAF was supported by University of Utah Developmental Biology Training Grant 5T32-HD07491. Work in SL’s laboratory was supported by the Swiss National Science Foundation (SNF_31003A_141093/1), the University of Zurich, the “Cells-in-Motion” Cluster of Excellence (EXC 1003-CiM), the Deutsche Forschungsgemeinschaft (DFG_LU 1398/2-1), and the University of Münster. Work in MMM’s laboratory was supported by National Institutes of Health (NIH) grant 1R01GM084011 and a March of Dimes Award 5-FY07-664.

---

**Supplemental Figure 1.**
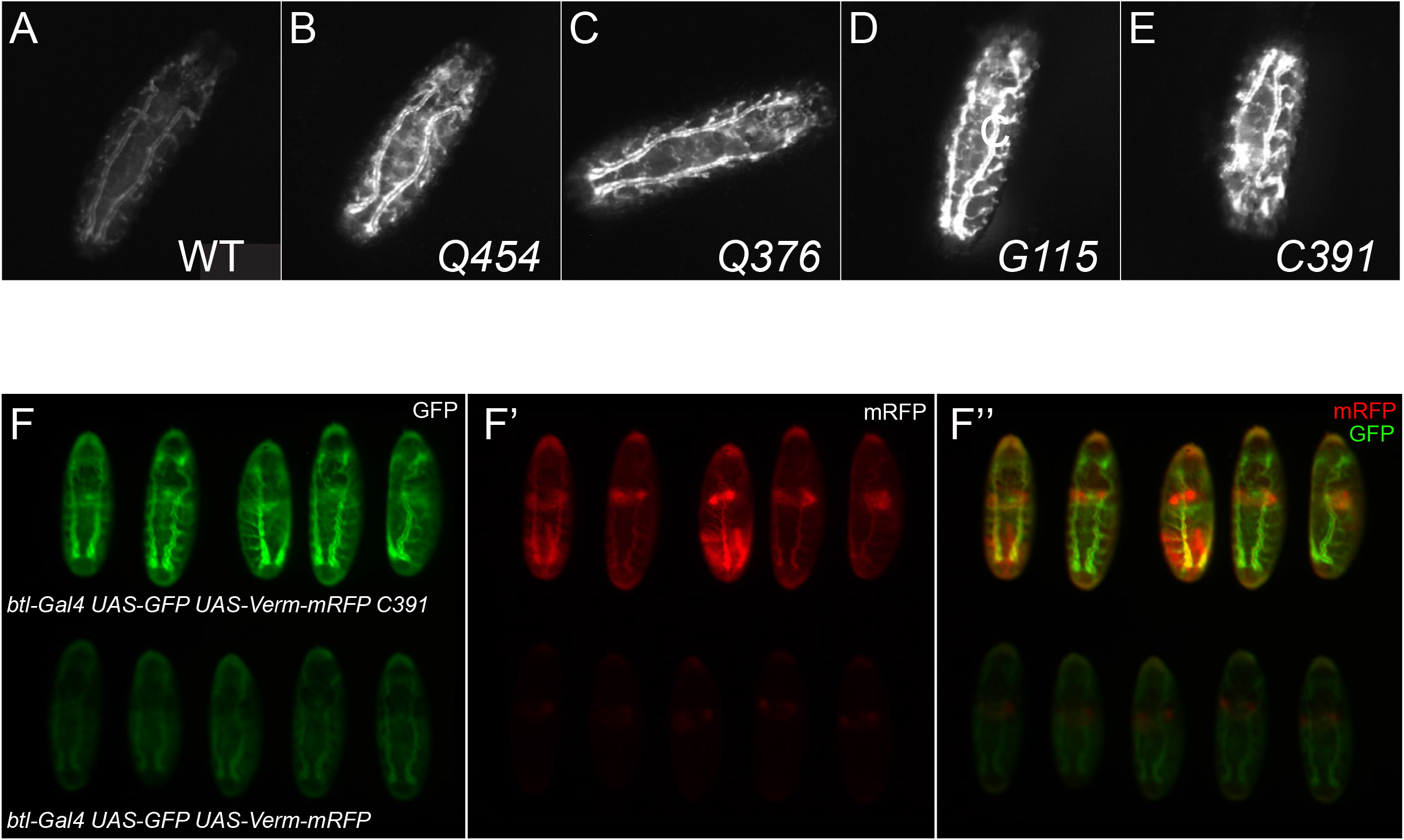
*Smg5* mutant alleles have enhanced fluorescence of NMD-sensitive reporters in first instar larval trachea. (**A-E)** NMD sensitive *eGFP::SV40 3′UTR* fluorescent reporter is expressed in larval trachea by the *btl-GAL4* driver in control (**A**) and *Smg5* mutant animals (**B-E**). **(F)** *UAS-GFP:SV40 3′UTR* and *UAS-VermmRFP:3′UTR* coexpressed in the embryonic tracheal system both show fluorescence enhancement (top row of embryos), compared to controls (bottom row of embryos).

**Supplemental Figure 2.**
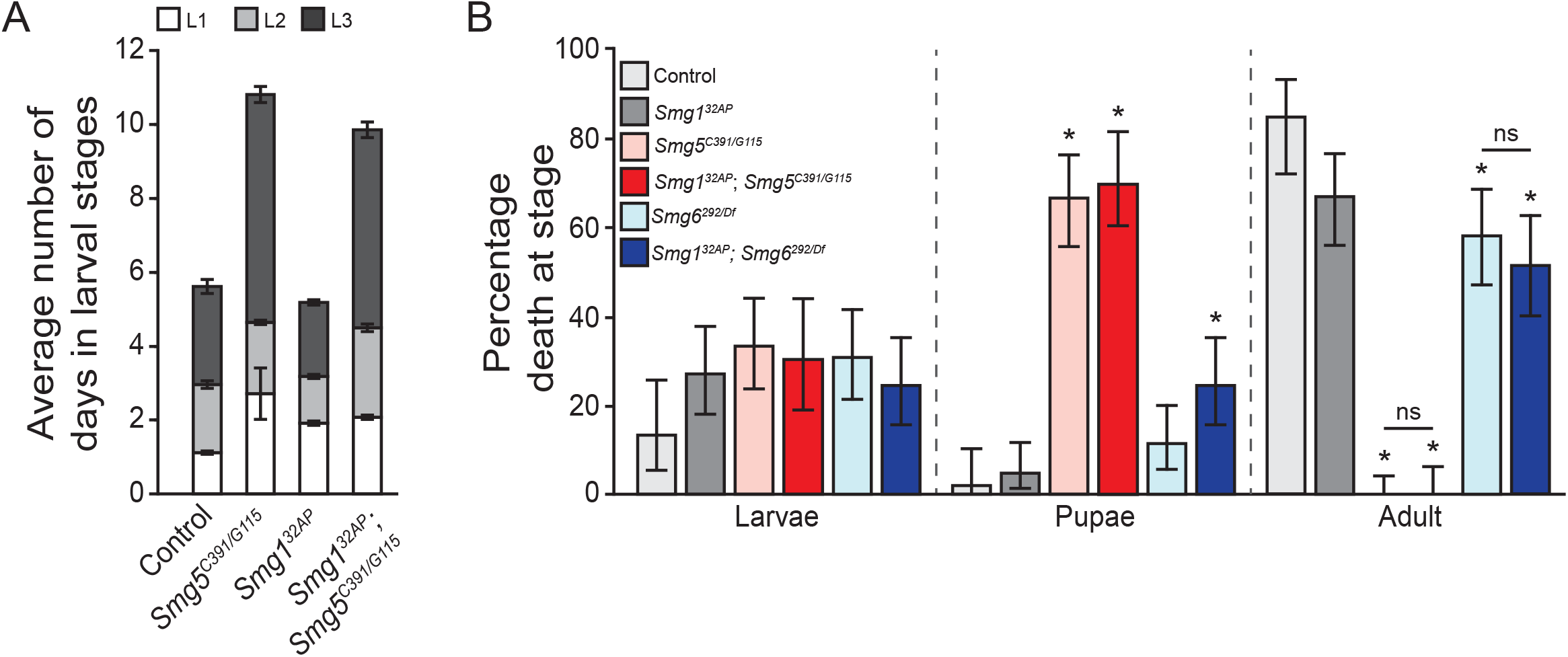
Lethal phase and developmental delays of single and double mutants for NMD genes. (**A**) Average number of days spent during larval stages of animals that entered pupariation in each indicated genotype. Error bars represent 2 SEM. (**B**) Percentage of animals that die during larval development, pupal development, or adulthood in indicated genotypes. Error represents 95% confidence interval of the binomial distribution. * indicates adjusted p-value < 0.05 compared to lethality of control condition at that stage.

**Supplemental Figure 3.**
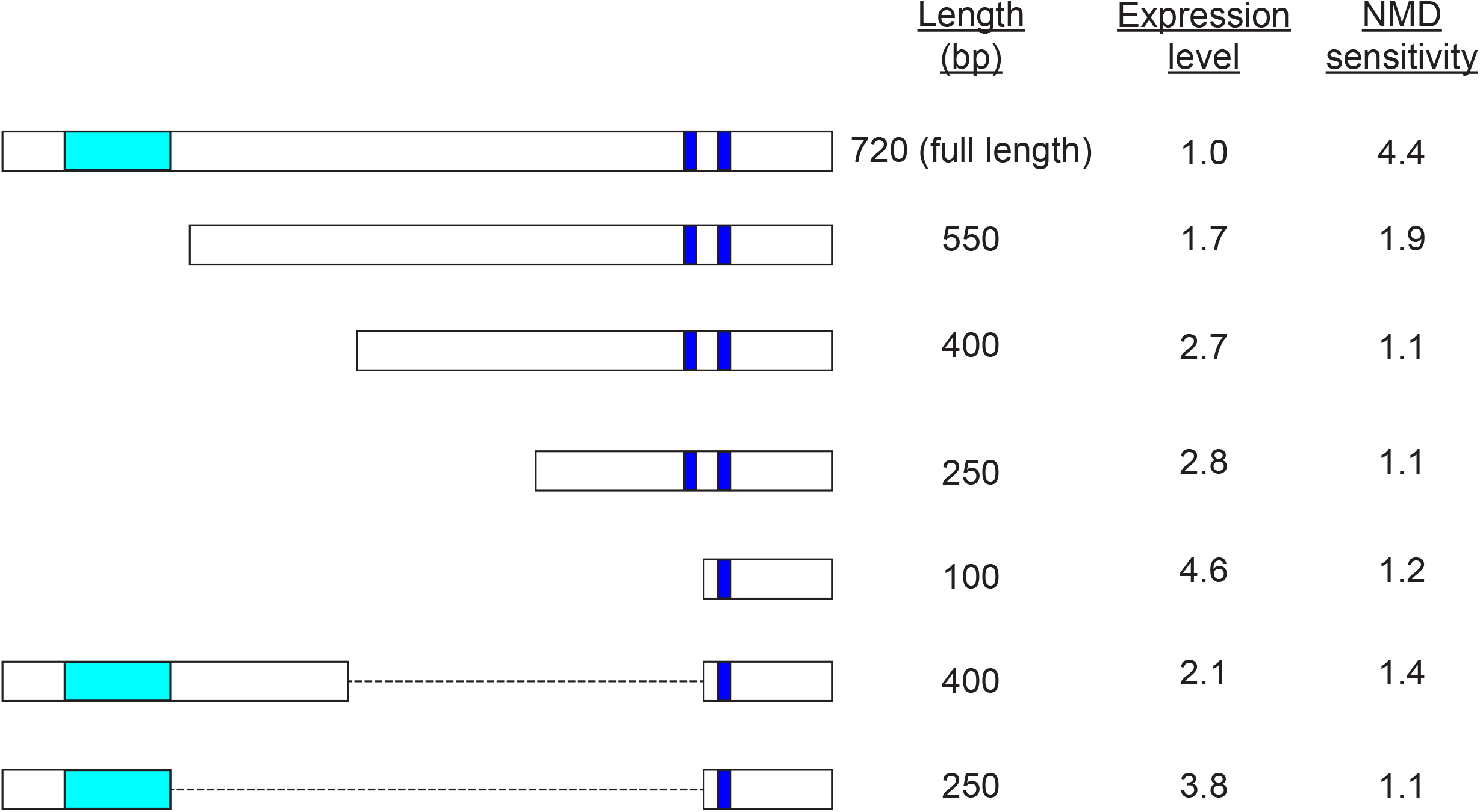
SV40 3′UTR deletion constructs tested for expression and NMD sensitivity. The full length 3′ UTR is shown in schematic and length given in base-pairs (bp). Cyan box, SV40 3′ UTR intron; blue boxes, predicted polyadenylation sites; dashed line, internal deletions. Expression level is the GFP signal observed when expressing the construct using an *e22c-GAL4* (epithelial) driver, normalized to the expression observed in the full-length construct. NMD sensitivity is the ratio of expression in an *Upf2^25G^* genetic background compared to a *Upf2^+^* genetic background (the higher the value the more sensitive the construct is to loss of NMD). For the experiments shown in Figure 3, the deletion used was the 100bp construct.

